# Network based conditional genome wide association analysis of human metabolomics

**DOI:** 10.1101/096982

**Authors:** Y. A. Tsepilov, S. Zh. Sharapov, O. O. Zaytseva, J. Krumsek, C. Prehn, J. Adamski, G. Kastenmüller, R. Wang-Sattler, K. Strauch, C. Gieger, Y. S. Aulchenko

## Abstract

**Background:** Genome-wide association studies (GWAS) have identified hundreds of loci influencing complex human traits, however, their biological mechanism of action remains mostly unknown. Recent accumulation of functional genomics (‘omics’) including metabolomics data opens up opportunities to provide a new insight into the functional role of specific changes in the genome. Functional genomic data are characterized by high dimensionality, presence of (strong) statistical dependencies between traits, and, potentially, complex genetic control. Therefore, analysis of such data asks for development of specific statistical genetic methods.

**Results:** We propose a network-based, conditional approach to evaluate the impact of genetic variants on omics phenotypes (conditional GWAS, cGWAS). For each trait of interest, based on biological network, we select a set of other traits to be used as covariates in GWAS. The network could be reconstructed either from biological pathway databases or directly from the data. We evaluated our approach using data from a population-based KORA study (n=1,784, 1.7 M SNPs) with measured metabolomics data (151 metabolites) and demonstrated that our approach allows for identification of up to five additional loci not detected by conventional GWAS. We show that this gain in power is achieved through increased precision of genetic effect estimates, and in presence of specific ‘contra-intuitive’ pleiotropic scenarios (when genetic and environmental sources of covariance are acting in opposite manner). We justify existence of such scenarios, and discuss possible applications of our method beyond metabolomics.

*Conclusions:* We demonstrate that in context of metabolomics network-based, conditional genome-wide association analysis is able to dramatically increase power of identification of loci with specific ‘contra-intuitive’ pleiotropic architecture. Our method has modest computational costs, can utilize summary level GWAS data, and is applicable to other omics data types. We anticipate that application of our method to new and existing data sets will facilitate progress in understanding genetic bases of control of molecular and complex phenotypes.

*Short abstract:* We propose a network-based, conditional approach for genome-wide analysis of multivariate omics phenotypes. Our methods can incorporate prior biological knowledge about biological pathways from external sources. We evaluated our approach using metabolomics data and demonstrated that our approach has bigger power and allows for identification of additional loci. We show that gain in power is achieved through increased precision of genetic effect estimates, and in presence of specific ‘contra-intuitive’ pleiotropic scenarios (when genetic and environmental sources of covariance are acting in opposite manner). We justify existence of such scenarios, and discuss possible applications of our method beyond metabolomics.

## Background

Genome-wide association studies (GWAS) is one of the most popular methods of identification of alleles that affect complex traits, including risk of common human diseases. In the past decade, GWAS allowed identification of thousands of loci, leading to a significant progress in understanding of genetic bases of control of complex human traits [1]. However, this had limited impact onto development of biomarkers and therapeutic agents, as most of the time the observation of association to a genomic region provides a starting point, but not yet a direct answer to the question of biological function affected by variation in the identified region. Recent accumulation of functional genomics data, which includes information on levels of gene expression (transcriptome), metabolites (metabolome), proteins (proteome) and glycosylation (glycome), could give a new insight into the functional role of specific changes in the genome [2,3]. Such data require special statistical methods for their analysis, because of their characteristically high dimensionality (ranging from few dozens to thousands and even to millions of measurements for each person), and presence of statistical dependencies reflecting biological relationships between individual omics components. Development of methods for omics data analysis is of current importance as the progress of molecular biology techniques continues and new types of functional genomic data become available.

Conventional univariate GWAS (uGWAS) ignore dependencies between different omics traits, which confounds biological interpretation of results and may lead to loss of statistical power. It was shown that utilizing multivariate phenotype representation increases statistical power, and leads to richer findings in the association tests compared to the univariate analysis [4–7]. Despite large number of methodological works, only few empirical multivariate GWAS have been published for humans. Among these which should be noted in relation to our work, Inouye et al. [8] performed multivariate GWAS of 130 NMR metabolites (grouped in 11 sets) in ~6600 individuals. The study demonstrated that multivariate analysis doubles the number of loci detected in this sample; among loci discovered via multivariate analysis seven were novel and did not appear before in other GWAS of related traits. While no replication of novel loci was performed in the original study, we compared results reported by Inouye et al. with recently published univariate GWAS of NMR metabolomics, which used sample size of up to 24,925 individuals [9]. We found that for three out of seven SNPs reported in the original work, p-value was < 5×10^−11^ for at least one metabolite. This provides empirical evidence for the value of multivariate methods in genomics of metabolic traits.

Here we propose a (knowledge-based) network-driven conditional genome-wide association analysis that exploits information from biologically related traits. To demonstrate our methodology, we performed proof-of-principle study directly comparing the power of univariate GWAS and the proposed method using metabolomics data (151 metabolites, Biocrates assay) from the KORA F4 study (n=1785).

## Results and Discussion

### Network-based conditional analysis of genetic associations

We start with theoretical justification and identification of specific scenarios under which adjustment for a biologically relevant covariate increases power of association analysis. Let us consider a trait of interest, y, covariate *c* and genotype g. Without loss of generality, assume that they are distributed with mean zero and standard deviation of one. Their joint distribution is specified by a set of three correlation coefficients, *𝜌*. Given specific parameter values, the value of “univariate” test statistic for association between *y* and *g* has the value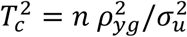, where *n* is the sample size and 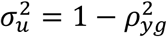 is the residual variance of *y*. For the conditional test, 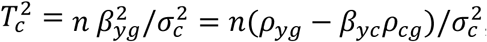, where *β* denote partial coefficients of regression from the conditional model and 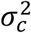 is the residual variance of *y*. Consequently, the log-ratio of these test statistics can be partitioned into two components

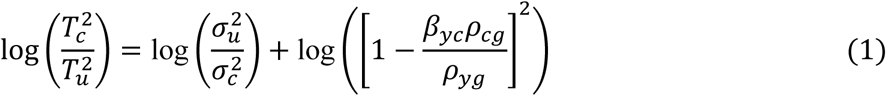

We shall call the first summand of (1) as ‘noise’ component and the second summand as the ‘pleiotropic’ component. Because the noise component 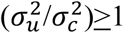 always, any possible reduction in the ratio between univariate and conditional test is determined by the sign and the magnitude of the term *β_yc_𝜌_cg_*/*𝜌_yg_*. When this product is negative, there is always increase in power of conditional analysis.

We can re-write *β_yc_𝜌_cg_*/*𝜌_yg_* as 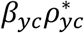, where 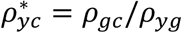 is a quantity which in a Mendelian randomization analysis is interpreted as the effect of the covariate on the trait free of non-genetic confounders [10]. Note that while 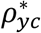 is reflecting the covariance between the trait and the covariate, which is induced by the effect of the genotype, *β_yc_* is related to ‘purely environmental’ sources of covariance between *y* and *c*. We can conclude that when genotype-induced and environmental correlations are consistent in sign, the product 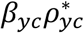 is positive and hence the contribution of the second term of (1) into relative power is negative. On the contrary, a ‘surprising’ product (where the sign is inconsistent and hence 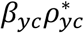 is negative) contribute positively to the relative power of conditional model.

In the context of complex polygenic traits, one expects that genetic and environmental correlations are consistent in sign. This is well reflected in animal breeding literature, and for a recent human example, one can see [11]. Under this scenario it would be desirable that *𝜌_cg_* (effect of genotype onto covariate) is very small, while *β_yc_* (which makes contribution into reduction of 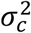 compared to 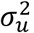 is large. However, in the context of specific locus affecting an activity of an enzyme involved in a biochemical reaction, the ‘surprising’ inconsistency between *β_yc_* and 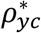 may be not so surprising. Indeed, consider an allele, which is associated with increased activity of an enzyme converting substrate A into product B. It is expected that A and B are positively correlated, and that the allele is in positive correlation with level of product B and in negative correlation with the substrate A. This is exactly a scenario which would lead to the positive value of the second term in (1), hence providing additional increase in power on the top of noise reduction.

We can readily extend the formula (1) to a case when *k* covariates are included in the conditional model. Denoting coefficients of correlation between *g* and covariate *i* as *𝜌_gi_* and partial coefficients of regression of *y* onto covariate *i* as *β_i_*, we have

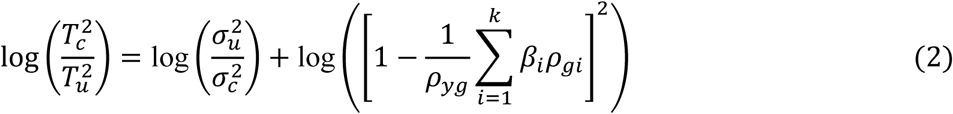

Above considerations allow us to hypothesize that a conditional GWAS (cGWAS), where covariates selected are biochemical, one-reaction-step neighbors of the target trait may provide increased power by exploiting both noise reduction and possible ‘surprising’ pleiotropy. In this work, we set off to empirically verify this hypothesis by investigating of human metabolomics data.

When proper covariates are selected, the methodology of cGWAS using individual-level data becomes rather trivial, and boils down to running a GWAS in which one jointly estimates the effect of an SNP and of specific covariates. The cGWAS method is less trivial in case one would like to exploit summary-level univariate GWAS data, for example these data which are available from previously published studies. Formulation of cGWAS on the level of summary GWAS statistics is possible, and we describe this method in Supplementary Note 1.

The question of selection of proper covariates is very important because it has direct consequences on the chances of finding the ‘surprising’ pleiotropic scenarios. In case biological/biochemical relations between the traits of interest are known and summarized in some database(s), this knowledge can be used directly by e.g. taking all direct neighbors as covariates. Alternatively, the network may be reconstructed in a hypothesis-free, empirical manner from the same or external data by e.g. using Gaussian graphical models (GGM) approach [12]; then some threshold may be applied to select the covariates.

### Comparison between cGWAS and uGWAS using human metabolomics data

We compared cGWAS and uGWAS methods using individual-level genetic and metabolomics data from KORA F4 study (1,784 individuals measured for 151 metabolite, Biocrates assay, and imputed at 1,717,498 SNPs).

First, we explored the potential of cGWAS where covariates were selected based on known biochemical network. Thus our analysis was restricted to a subset of 105 metabolites for which the one-reaction-step immediate biochemical neighbors were available [12]. This biochemical network incorporates only lipid metabolites, and pathway reactions cover two groups of pathways: (1) Fatty acid biosynthesis reactions which apply to the metabolite classes lyso-PC, diacyl-PC, acyl-alkyl-PC and sphingomyelins; (2) β-oxidation reactions representing fatty acid degradation to model reactions between the acyl-carnitines. The β-oxidation model consists of a linear chain of C2 degradation steps (C10-C8-C6 etc.). Number of covariates varied from one to four with mean of 2.48 and median 2.

**Table 1** shows 11 loci which were significant in either cGWAS or uGWAS analysis and fall into known regions (see Supplementary Note 2). Of these, ten loci were identifiable by cGWAS and nine were identifiable by uGWAS. Compared to uGWAS, one locus (*ETFDH*) was lost, but two additional loci were identified (*ACSL1* for PC ae C42:5, and *PKD2L1* for lysoPC a C16:1). It is interesting to note that for *ACSL1* (SNP rs4862429 effect onto PC ae C42:5, with cGWAS p=7e-11), the uGWAS p-value was 0.7. This is expected under the model of ‘surprising’ pleiotropy.

**Table 1.**
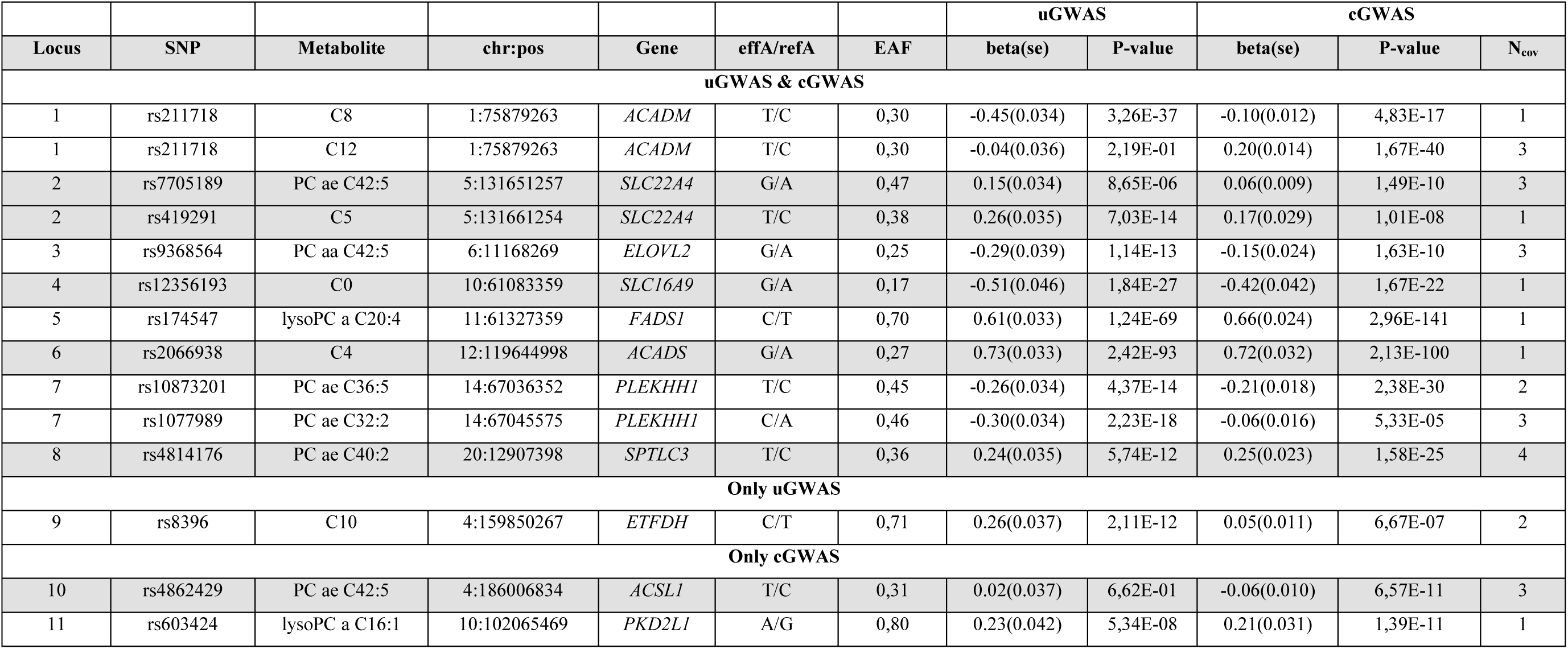
Eleven loci found by cGWAS and uGWAS on metabolites for which at least one one-reaction-step neighbor was available. Best SNP - Metabolite pair is shown for each locus. *chr:pos* corresponds to the physical position of SNP; EAF - effect allele frequency, beta(se) - estimated effect and standard error of the SNP; effA/refA - effect/reference alleles; P-value - p-value of the additive model; *Gene* - the most probable (according to DEPICT) associated gene in the region; *N_cov_* – number of covariates used in cGWAS.

To test whether use of cGWAS increases average power of association analysis, we contrasted the average of cGWAS and uGWAS maximal chi-squared test statistics for loci from Table 1. The ratio of average maximal test statistic between cGWAS and uGWAS was 1.59. However, the Wilcoxon paired sample test contrasting the best cGWAS vs. the best uGWAS values of chi-squared test statistic, was only marginally significant (p=0.067).

For the SNPs listed in **Table 1**, we applied formula (2) to partition the log-ratio of the cGWAS and uGWAS test statistics into ‘noise’ and ‘pleiotropic’ components. **Figure 1** shows that the trend in the ratio is mainly determined by the second (‘pleiotropic’) summand. One can see that, with the exception of locus *SLC22A4,* SNP-trait pairs for which cGWAS had increased power are these where the second term of (1) is positive or close to zero. In contrast, the SNP-trait combinations which were lost in cGWAS, had strong negative contribution from the ‘pleiotropic’ term of (2).

**Figure 1.**
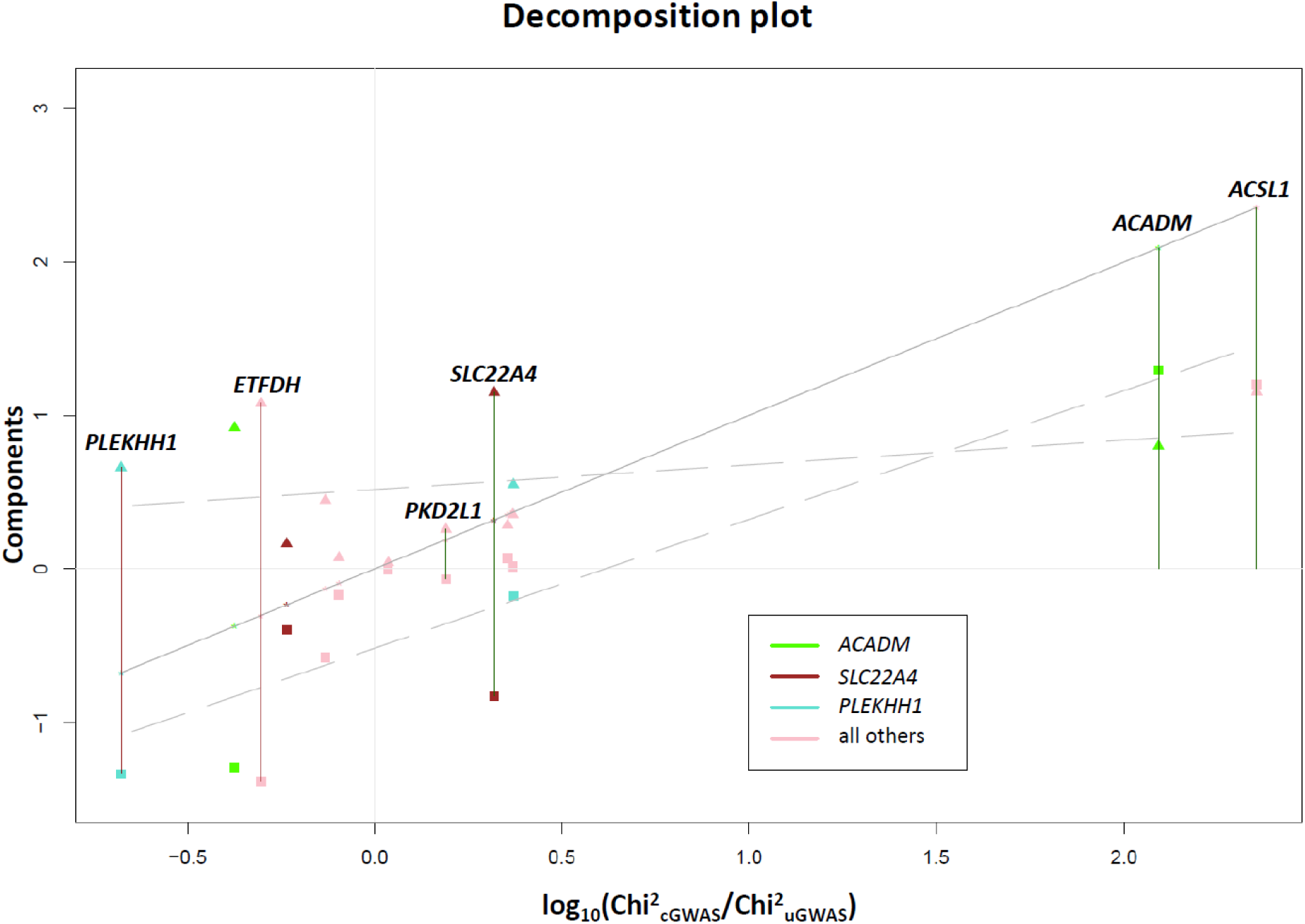
Decomposition of Chi-squared ratio for cGWAS and uGWAS method into pleiotropic and noise components. The stars correspond to the sum of components that is Chi-squared ratio (y=x line). Pleiotropic component is represented by squares, noise component – by triangles. Dashed lines correspond to regression lines for the two component. Dark green vertical lines indicate SNP-trait combinations that were significant in cGWAS and not significant in uGWAS; dark red line indicates the SNP-trait combinations which was significant in uGWAS only.

It is interesting to investigate the variance-covariance structure of loci with positive and negative pleiotropic term. We selected two loci where the pleiotropic component’s contribution to power was positive (rs174547 at *FADS1* locus) and negative (rs8396 at *ETFDH*). We show corresponding correlations between SNP and trait and covariates involved, together with partial coefficients from conditional regression of the trait onto SNP and covariates in Figure 2. For *FADS1* locus (Figure 2A), the correlation between SNP and the trait (lysoPC a C20:4) and the covariate (lysoPC a C20:3) are in opposite directions, while the trait and the covariate are positively correlated (both based on correlation and partial correlation). As a consequence, we can see that the value of partial regression coefficient between the SNP and lysoPC a C20:4, conditional on lysoPC a C20:3 is greater than coefficient of regression without covariates. This makes biological sense as *FADS1* is coding the fatty acid desaturase enzyme, while these two traits differ from each other by one double bond. It appears that this case suits perfectly the biochemical scenario under which we expect increased power of conditional analysis.

**Figure 2.**
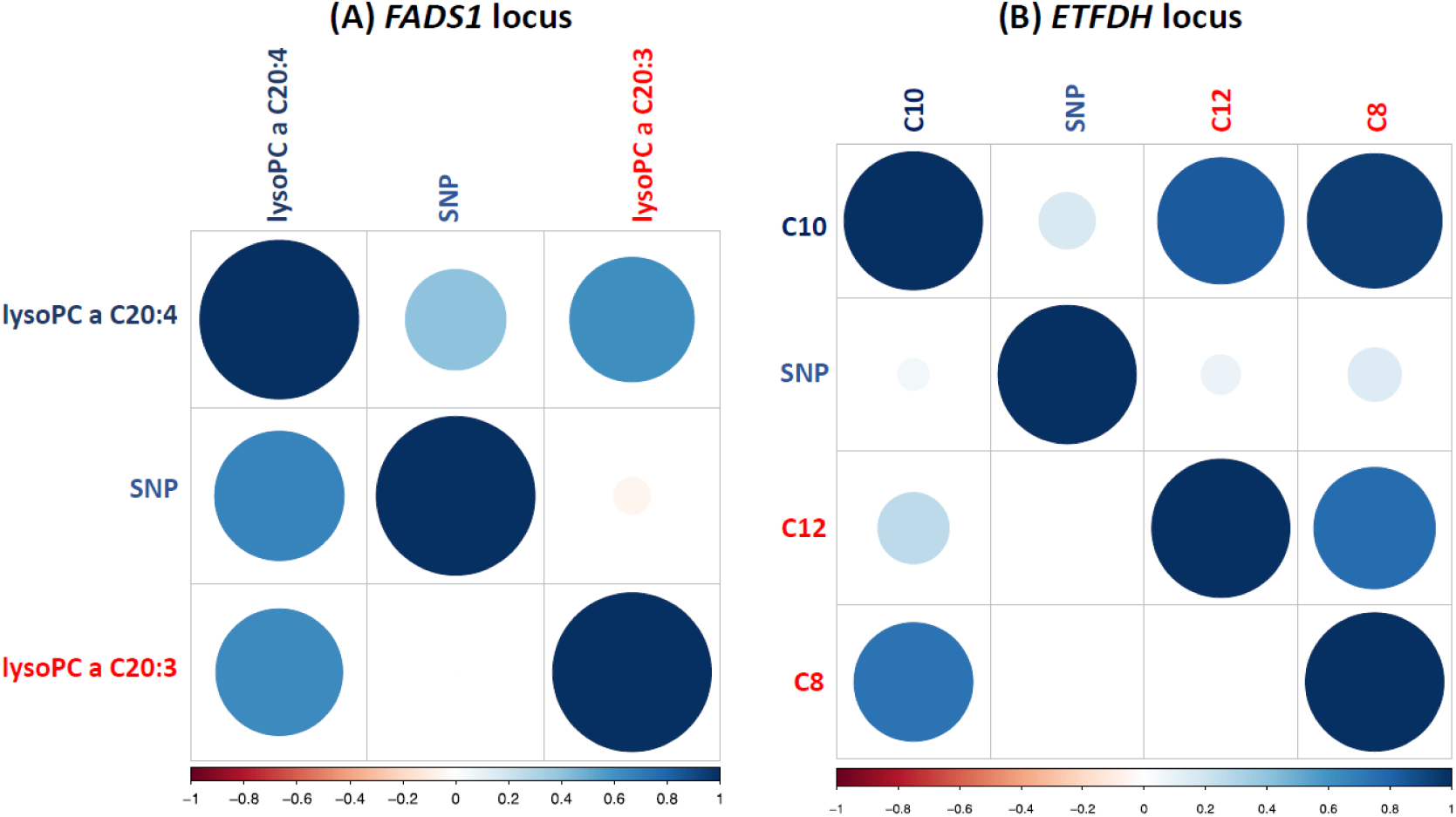
Correlations (above diagonal) and partial coefficients of regression of the trait of interest (below diagonal) for *FADS1* and *ETFDH* loci, representing scenarios in which pleiotropic term of (2) is strongly positive and negative respectively.

In the second example (Figure 2B, *ETFDH*), we observe that conditional regression of C10 onto rs8396 and two covariates (C8 and C12, medium-chain acylcarnitines) leads to smaller SNP coefficient compared to unconditional regression; this happens because all terms of 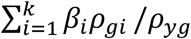 are positive. The *ETFDH* gene, prioritised as the best candidate by DEPICT (FDR<5%), encodes for electron transfer flavoprotein dehydrogenase that is involved into fatty acid oxidation in the mitochondria. During this process the acyl group is transferred from long chain acylcarnitines to form long-chain acetyl-CoA, which is then catabolized. ETF dehydrogenase takes part in the catabolic process by transferring electrons from Acyl-CoA dehydrogenase into the oxidative phosphorylation pathway. Thus, the ETFDH gene should act on all kinds of long-chain acylcarnitines in the same direction and we can expect that pleotropic influence of this gene onto the acylcarnitines in our example (C8, C10, C12) will be unidirectional. Presence unidirectional genetic effects and positive correlations between these acylcarnitines makes second term of equation (2) negative, which leads to the decreased power of genetic association analysis.

Above analysis provide a real-life example that use of biochemical neighbors to adjust genetic association analysis of target trait allows for (sometimes very sharp) increase of power for the genetic variants which act in ‘surprising’ pleiotropic manner; our analysis also suggests that cGWAS may increase GWAS power on average, although this increase is not uniform and heavily depends on pleiotropic relations between involved locus and the traits.

While use of known biochemical network for covariate selection has many attractive properties, it may be somewhat unpractical, because our biochemical knowledge is yet fragmented. Therefore, next we have investigated the potential of cGWAS method where covariates are selected using data-driven approach. The metabolites network was reconstructed using Gaussian Graphical Models based on partial correlations. For a target metabolite, covariates were selected based on significant partial correlations. For that, we have chosen threshold proposed previously in [12]: p-value≤(0.01/Number of calculated partial correlations), which corresponds to a cut-off p-value≤8.83×10^−7^. The network used in our analysis is presented in **Supplementary Figure 1.** For the clarity of notation, hereafter we will call cGWAS using known biochemical network as BN-cGWAS, and cGWAS which is based on GGM selection of covariates as GGM-cGWAS.

To contrast GGM-cGWAS and BN-cGWAS, we first used the same set of metabolites which was utilized by BN-cGWAS to run GGM-cGWAS. The results are presented in **Supplementary Table 1**. We found 16 SNP-trait pairs clustered to 10 loci that could be detected by GGM-cGWAS or BN-cGWAS. The number of covariates included into GGM-cGWAS analysis, was larger (from 2 to 18, with mean of 8.5) than that in BN-cGWAS. Therefore, we expected that GGM-cGWAS may gain relative power compared to BN-cGWAS because of noise reduction (term 1 of equation(2)); however, we it may also be expected that GGM-cGWAS may lose power because of less likely occurrence of ‘surprise’ pleiotropy (term 2 of equation (2)).

For the best SNP-trait pairs detected by GGM-cGWAS or BN-cGWAS, we computed the components of equation (2) and contrasted them using Wilcoxon paired samples test. The noise component of (2) was always greater for GGM-cGWAS (mean difference of 0.66, p=3×10^−5^). For GGM-cGWAS, the second ‘pleiotropic’ component of equation (2) was on average smaller than that for the BN-cGWAS (mean difference −0.54, p=0.013); still, for three GGM-cGWAS SNP-trait pairs out of 16 the pleiotropic component was positive. Average Chi-squared statistics was 33% smaller for GGM-cGWAS that for BN-cGWAS indicating average loss of power (although this loss was not significant, Wilcoxon paired test p=0.5), but at the same time it still was 22% bigger than uGWAS (Wilcoxon paired test p=0.8). We conclude that while GGM-cGWAS is in a way imperfect proxy to use of real biochemical network, it may still have increased power because of even further reduced target trait residual variance, and some potential to detect ‘surprising’pleiotropy.

To explore the potential of cGWAS under realistic conditions to a full extent, we analyzed all 151 available metabolites using GGM-cGWAS and contrasted the results to uGWAS (Table 2 and **Supplementary Figure 2)**. In total, uGWAS was able to detect 15 loci at genome-wide significance level defined as p≤5×10^−8^/151 = 3.3×10^−10^. Applying GGM-cGWAS, we identified 19 significant loci at the same threshold. Expectedly, we observed that compared to uGWAS the precision of genetic effect estimation increased (Table2, Supplementary Figure 3). The overlap between uGWAS and GGM-cGWAS findings was 14 loci, with GGM-cGWAS losing one locus (for C5:1-DC at rs2943644), but identifying five new loci not identified by uGWAS. Three of the five new loci were affecting amino acids, and two – acylcarnitines. Note that loci identified by BN-cGWAS (covariates selected via biochemical network) are a subset of 19 loci identified by GGM-cGWAS.

We have investigated the literature results available for the loci described in Table 2 (see Supplementary Note 2 for details). From 20 loci we report in this study, 15 were genome-wide significant in recent large (n=7,478) meta-analysis of Biocrates metabolomics data by Draisma *et al.* [13]. For 11 of 15 loci, we observed significant association for exactly the same SNP-metabolite pair. However, not all metabolites analyzed in this study were analyzed by Draisma *et al.* [13]; still, for the residual three loci the top association was with a metabolite within the same class as in our study and one from different lipid classes (**see Supplementary Table 2**). For the other five loci, which did not show significant association in work of Draisma *et al.* [13], we have checked if these were significant and replicated in work of Tsepilov et al. [14]. It should be noted though that in work [14], the same KORA F4 data set was used as discovery, and the analysis concerned the ratios of metabolites. Out of five loci, two were significant and replicated in [14], and in all two cases, the metabolite analyzed in this work was the part of the ratio analyzed by Tsepilov et al. One of five was published before for the same trait in other studies [15,16]. We did not find previous evidence for association with metabolites for rs2943644 (*LOC646736*) and rs17112944 (*LOC728755*). Therefore, we are inclined to consider observed associations with rs17112944 and rs2943644 as potential false positives; these two loci are excluded from further consideration.

**Table 2.** Twenty loci found by cGWAS and uGWAS approaches. Best SNP - Metabolite pair is shown for each locus. *chr:pos* corresponds to the physical position of SNP; EAF - effect allele frequency, beta(se) - estimated effect and standard error of SNP; effA/refA - effect/reference alleles; P- value - p-value of the additive model; *Gene* - the most probable (according to DEPICT) associated gene in the region; *N_cov_* – number of covariates for cGWAS.

|  |  |  |  |  |  |  | uGWAS |  | cGWAS |  |  |
| --- | --- | --- | --- | --- | --- | --- | --- | --- | --- | --- | --- |
| Locus | SNP | Metabolite | chr:pos | Gene | effA/refA | EAF | beta(se) | P-value | beta(se) | P-value | N<sub>cov</sub> |
| uGWAS & cGWAS |  |  |  |  |  |  |  |  |  |  |  |
| 1 | rs211718 | C6 (C4:1-DC) | 1:75,879,263 | ACADM | T/C | 0.30 | -0.48(0.034) | 4.64E-42 | -0.13(0.017) | 2.00E-13 | 7 |
| 1 | rs7552404 | C6 (C4:1-DC) | 1:75,908,534 | ACADM | G/A | 0.30 | -0.48(0.034) | 3.10E-42 | -0.12(0.017) | 3.25E-13 | 7 |
| 2 | rs483180 | Ser | 1:120,069,028 | PHGDH | G/C | 0.30 | -0.24(0.037) | 3.34E-11 | -0.24(0.028) | 1.50E-17 | 2 |
| 2 | rs477992 | Ser | 1:120,059,099 | PHGDH | A/G | 0.70 | 0.24(0.037) | 5.15E-11 | 0.24(0.028) | 5.82E-18 | 2 |
| 3 | rs2286963 | C9 | 2:210,768,295 | ACADL | G/T | 0.63 | -0.49(0.032) | 1.10E-49 | -0.48(0.027) | 1.48E-67 | 3 |
| 4 | rs8396 | C10 | 4:159,850,267 | ETFDH | C/T | 0.71 | 0.26(0.037) | 2.02E-12 | 0.04(0.010) | 1.49E-05 | 8 |
| 4 | rs8396 | C7-DC | 4:159,850,267 | ETFDH | C/T | 0.71 | -0.09(0.037) | 1.67E-02 | -0.13(0.020) | 3.29E-11 | 8 |
| 5 | rs419291 | C5 | 5:131,661,254 | SLC22A4 | T/C | 0.38 | 0.26(0.035) | 7.03E-14 | 0.17(0.026) | 2.28E-10 | 3 |
| 5 | rs270613 | C5 | 5:131,668,482 | SLC22A4 | A/G | 0.61 | -0.26(0.035) | 7.93E-14 | -0.17(0.026) | 8.48E-11 | 3 |
| 6 | rs9393903 | PC aa C42:5 | 6:11,150,895 | ELOVL2 | A/G | 0.75 | 0.29(0.039) | 2.19E-13 | 0.18(0.020) | 4.51E-19 | 6 |
| 6 | rs9368564 | PC aa C42:5 | 6:11,168,269 | ELOVL2 | G/A | 0.25 | -0.29(0.039) | 1.14E-13 | -0.19(0.021) | 7.84E-19 | 6 |
| 7 | rs816411 | Ser | 7:56,138,983 | PHKG1 | C/T | 0.51 | -0.22(0.034) | 2.15E-10 | -0.19(0.026) | 5.16E-13 | 2 |
| 7 | rs1894832 | Ser | 7:56,144,740 | PHKG1 | C/T | 0.51 | 0.21(0.034) | 3.23E-10 | 0.19(0.026) | 1.69E-13 | 2 |
| 8 | rs12356193 | C0 | 10:61,083,359 | SLC16A9 | G/A | 0.17 | -0.51(0.046) | 1.84E-27 | -0.27(0.034) | 9.72E-16 | 3 |
| 9 | rs174547 | lysoPC a C20:4 | 11:61,327,359 | FADS1 | C/T | 0.70 | 0.61(0.033) | 1.44E-69 | 0.07(0.011) | 1.41E-10 | 9 |
| 9 | rs174556 | PC ae C44:4 | 11:61,337,211 | FADS1 | T/C | 0.27 | 0.09(0.038) | 1.55E-02 | 0.21(0.014) | 3.16E-46 | 3 |
| 10 | rs2066938 | C4 | 12:119,644,998 | ACADS | G/A | 0.27 | 0.73(0.033) | 5.87E-94 | 0.71(0.025) | 1.31E-151 | 2 |
| 11 | rs12879147 | PC aa C28:1 | 14:63,297,349 | SYNE2 | A/G | 0.85 | -0.46(0.050) | 2.07E-19 | -0.12(0.019) | 5.94E-11 | 14 |
| 11 | rs17101394 | SM(OH) C14:1 | 14:63,302,139 | *SYNE2* | A/G | 0.83 | -0.32(0.050) | 1.00E-10 | -0.10(0.011) | 1.17E-17 | 7 |
| 12 | rs1077989 | PC ae C36:5 | 14:67,045,575 | *PLEKHH1* | C/A | 0.46 | -0.26(0.034) | 3.42E-14 | -0.08(0.010) | 8.25E-16 | 10 |
| 12 | rs1077989 | PC ae C32:2 | 14:67,045,575 | *PLEKHH1* | C/A | 0.46 | -0.30(0.034) | 2.23E-18 | -0.05(0.016) | 1.31E-03 | 6 |
| 13 | rs4814176 | SM(OH) C22:1 | 20:12,907,398 | *SPTLC3* | T/C | 0.36 | 0.03(0.035) | 4.51E-01 | -0.07(0.009) | 1.10E-16 | 10 |
| 13 | rs4814176 | SM(OH) C24:1 | 20:12,907,398 | *SPTLC3* | T/C | 0.36 | 0.24(0.035) | 5.40E-12 | 0.09(0.013) | 3.04E-11 | 9 |
| 14 | rs5746636 | Pro | 22:17,276,301 | *PRODH* | T/G | 0.24 | -0.31(0.039) | 3.00E-15 | -0.32(0.034) | 1.91E-20 | 2 |
| Only uGWAS |  |  |  |  |  |  |  |  |  |  |  |
| 15 | rs2943644 | C5:1-DC | 2:226,754,586 | *LOC646736* | C/T | 0.68 | 0.32(0.042) | 5.14E-14 | 0.09(0.022) | 3.58E-05 | 5 |
| Only cGWAS |  |  |  |  |  |  |  |  |  |  |  |
| 16 | rs1374804 | Gly | 3:127,391,188 | *ALDH1L1* | A/G | 0.64 | 0.20(0.036) | 1.88E-08 | 0.21(0.030) | 8.08E-13 | 3 |
| 17 | rs4862429 | PC ae C42:5 | 4:186,006,834 | *ACSL1* | T/C | 0.31 | 0.02(0.037) | 6.62E-01 | -0.06(0.008) | 1.25E-12 | 8 |
| 18 | rs603424 | C16:1 | 10:102,065,469 | *PKD2L1* | A/G | 0.80 | 0.16(0.042) | 9.51E-05 | 0.14(0.018) | 1.32E-13 | 9 |
| 19 | rs2657879 | Gln | 12:55,151,605 | *GLS2* | G/A | 0.21 | -0.24(0.042) | 2.82E-08 | -0.27(0.031) | 9.37E-18 | 5 |
| 20 | rs17112944 | C6:1 | 14:27,179,297 | *LOC728755* | A/G | 0.90 | -0.28(0.059) | 2.09E-06 | -0.21(0.032) | 1.38E-10 | 9 |

## Conclusions

We have developed a new approach for network-based conditional genome-wide association study for metabolomics data (conditional GWAS, cGWAS). For each metabolite trait, we select a set of other metabolites, to be used as covariates in GWAS. The selection of covariates could be done in a mechanistic way, e.g. based on known biological relations between traits of interest; or in a data-driven way, e.g. based on partial correlations. The method has modest computational costs and can exploit either individual- or summary-level GWAS data. It has a potential to increase the power of genetic association analysis because of reduced noise and ability to detect specific pleiotropic scenarios, hardly detectable via standard single-trait GWAS.

We have applied cGWAS approach to analysis of 151 metabolomics traits (Biocrates panel) in large (n=1,784) population-based KORA cohort. While conventional uGWAS identified 15 loci in this data set, cGWAS was able to identify up to 5 additional loci. At the same time, we have observed that for some loci the power of cGWAS was decreased. We found that in cGWAS power is always gained because of increased precision of genetic effect estimation, but it may be decreased or increased in presence of specific pleiotropic association scenarios.

We show that conditional analysis has especially high power under scenarios when locus-specific genotypic and environmental sources of covariance between the trait and its covariates are ‘surprising’ (acting in opposite direction). This type of pleiotropy is not unexpected for metabolic traits, and we provide an empirical demonstration of existence of such scenarios in this work. This is further demonstrated by the fact that the power gain from the pleiotropic component was higher when we used a mechanistic way of covariate selection (one-reaction-step neighbors from a biochemical network), as opposed to data-driven network (based on Gaussian Graphical Model). We may expect that with increased knowledge of biological networks the mechanistic way of covariate selection may become preferable.

However, when genotypic and environmental sources of covariance are consistent, cGWAS may lose power even compared with standard GWAS without biological covariates. One may argue that a joint analysis testing effects of genotype on the set of traits simultaneously may be a better solution, which maintains power across wide range of scenarios. While we are not arguing with this viewpoint, we must emphasize one aspect which makes conditional analysis attractive; namely, better interpretability of the obtained results in terms of effect of genotype on specific trait. The latter may be important in the next step when we may try to relate obtained results with these obtained previously for other traits in other GWAS, e.g. using methods described by [17–19].

Presence of highly correlated traits and different pleiotropic scenarios are not unique for metabolomics. Therefore, we expect that cGWAS may be a powerful approach for investigation of other omics traits. Low computational costs and possibility of analysis based on summary-level data makes cGWAS a promising approach to investigate new and re-analyze existing omics data sets in order to provide deeper understanding of functional genomics.

## Materials and Methods

### KORA study

The KORA cohort (Cooperative Health Research in the region of Augsburg) are population-based studies from the region of Augsburg in Southern Germany [20]. The KORA F4 is the follow-up survey (from 2006 to 2008) of the base line survey KORA S4 that was conducted from 1999 to 2001. All study protocols were approved by the ethics committee of the Bavarian Medical Chamber (Bayerische Landesärztekammer), and all participants gave written informed consent.

Concentrations of 163 metabolites were quantified in 3,061 serum samples of KORA F4 participants using flow injection electrospray ionization tandem mass spectrometry and the Absolute*IDQ*^TM^ p150 Kit (BIOCRATES Life Sciences AG, Innsbruck, Austria) [21]. After quality control 151 metabolite measurements were used in analysis. Details of the methods and quality control of the metabolite measurements and details of the metabolite nomenclatures were given previously [21]. Metabolite nomenclatures could be found in Supplementary Table 3.

Genotyping was performed with the Affymetrix 6.0 SNP array (534,174 SNP markers after quality control) with further imputation using HapMap2 (release 22) as reference panel resulting in a total of 1,717,498 SNPs (details given in Kolz *et al.* 2009 [22]). For 1,785 individuals both metabolite concentrations and genotypes were available in the KORA F4 study.

### Statistical analysis

Calculation of partial correlations and their p-values were performed using “ppcor” [23] R library. Graphical representations were made by “ggm” [24] R library. Similar to previous work [12], we considered partial correlation coefficient as significant if correlation’s p-value was less than 01/(151*150/2) (8.83×10^−7^).

For the GWAS analysis we used OmicABEL software [25]. All traits were first adjusted for sex, age and batch effect, and then residuals were transformed using inverse-normal transformation [26] prior to GWAS. The genotypes from KORA F4 were used. Only SNPs that had a call rate ≥ 0.95, R^2^ ≥ 0.3, Hardy–Weinberg equilibrium (HWE) p ≥ 10^−6^ and MAF ≥ 0.1 (1,717,498 SNPs in total) were considered in analysis. The genomic control method was applied to correct for a possible inflation of the test statistics. Lambda for all traits was between 1.00 and 1.03. To define independent loci, we have selected all genome-wide significant SNP-trait pairs, and identified the groups which were separated by >500kb. For regions of association, the most associated SNP-trait pair (as indicated by the lowest p-value) was selected to represent this locus. cGWAS and uGWAS results were considered to come from different loci if top SNPs were separated by >500kb. The threshold for GWAS analysis for 151 traits was p-value=5e-8/151=3.31×10^−10^.

When partitioning log(cGWAS/uGWAS) test statistic into noise and pleiotropic components (equation (2), Figure 1), we used all known loci that were significant in either cGWAS or uGWAS analyses. If locus included two SNP-trait pairs and traits were different we included both. If locus consisted two SNP-trait pairs and traits were the same, we included the one with lowest uGWAS p-value. When comparing the pleiotropic and noise components, the Wilcoxon paired samples test was used to perform statistical significance testing. For contrasting values of chi-squared test statistics, we employed similar procedure, with the exception that if results from specific analysis for specific locus were not genome-wide significant, for this method we have selected the maximal chi-squared test statistic from the +/−500kb region centered at the top association detected by the alternative method.

### *In silico* functional annotation

We conducted functional annotation of the novel discoveries. For prioritizing genes in associated regions, gene set enrichment and tissue/cell type enrichment analyses, we used the DEPICT software v. 140721 [27] with following parameters: flag_loci = 1; flag_genes = 1; flag_genesets = 1; flag_tissues = 1; param_ncores = 2, and further manual annotation (h37 assembly). All 27 SNPs clustered in 20 loci found by cGWAS and uGWAS (Table 2) were included into analysis. If several genes were proposed for a SNP by DEPICT we selected the gene with the lowest nominal DEPICT P-value. In most of the cases the results of manual annotation matched with the results of DEPICT annotation (see Supplementary Note 2). Additionally, we have looked up each SNP using the Phenoscanner [28] database to check whether it was previously reported to be associated with metabolic traits with p-value lower than 5×10^−8^ and proxy r^2^ =0.7.

## Additional files

Supplementary Note 1 – cGWAS using summary level data

Supplementary Note 2 – Literature search of loci identified by cGWAS and uGWAS

Supplementary Tables

ST 1 – BD-GWAS and GGM-GWAS for 105 metabolites
ST 2 – GGM-cGWAS and uGWAS for 151 metabolites
ST 3 - List of metabolites measured with the AbsoluteIDQ^®^ p150 Kit

Supplementary Figures

SF 1 – Partial correlations network
SF 2 – Manhattan plots for cGWAS and uGWAS for 151 metabolites
SF 3 – Comparison of effect estimates and their standard errors for SNPs from Table 2

## Abbreviations

GWAS: genome wide association study
cGWAS: conditional GWAS
uGWAS: univariate GWAS (trait-by-trait)
BN-cGWAS: cGWAS based on biochemical networks
GGM-cGWAS: cGWAS based on partial correlations network

## Acknowledgements

We would like to thank Athina Spilopoulou and Felix Agakov for fruitful discussion. We thank Alexander Zlobin and Alexander Grishenko for help with arranging tables and figures for the manuscript. We would like to express our gratitude to Sophie Molnos for help with management of the data.

## Authors contribution

YT, CG, YA planned and supervised the study; PC, CP and JA, KG, RW-S collected data, CG, KS contributed data for analysis; YT, OZ, SS performed data analysis; YT, YA, CG, OZ, JK, KS discussed and interpreted the results; YT, OZ, CG, YA wrote the manuscript. All authors have corrected and approved the final version of the manuscript.

